# A fluorescent biosensor for the visualization of agmatine

**DOI:** 10.1101/2024.07.27.605435

**Authors:** Pascal Kröger, Andre C. Stiel, Birthe Stüven, Sooruban Shanmugaratnam, Susanne Schoch, Dagmar Wachten, Birte Höcker

## Abstract

Agmatine is known to regulate various neurotransmitter systems, yet the underlying molecular mechanisms remain elusive. To visualize agmatine dynamics in cellular networks and thereby unravel its physiological functions, we developed a genetically encoded fluorescent agmatine biosensor (AGMsen) based on the periplasmic putrescine binding protein PotF. We first analyzed the agmatine binding properties of the PotF receptor module based on its crystal structure and introduced mutations into the binding pocket to favor agmatine binding over other biogenic amines. Validation of AGMsen functionality across different cell types, including primary neuronal cultures, demonstrates its ability of real-time agmatine visualization. Thus, the sensor can serve as a valuable tool to advance our understanding of agmatine distribution and dynamics, and shed light on their effect on neuronal functions *in vivo*.

## Introduction

Neurotransmitters are key regulators of brain function, and their dysregulation leads to several neuropathologies^1–3^. The use of optical imaging to reveal spatio-temporal neurotransmitter distributions largely contributes to understanding neuronal function and to the development of new concepts for therapies to treat neuropathologies. Prime examples are genetically encoded biosensors that allow to visualize neurotransmitters like dopamine, glutamate, GABA, and serotonin in different organisms^4–8^. A neurotransmitter that is yet at the advent of its research is agmatine (AGM), the decarboxylated form of the amino acid arginine. AGM can be taken up by axon terminals and is localized in synaptic vesicles, from which it can be released in a calcium-dependent manner^9,10^. This is in line with the proposed neurotransmitter-like function of AGM since it shows an influence on multiple molecular targets that include neurotransmitter systems such as nicotinic, imidazoline I_1_ and I_2_, α_2_-adrenergic, glutamate NMDAR, and serotonin 5-HT2A and 5HT-3 receptors^11^. Even with this multitude of different targets, agmatine is still referred to as a neuromodulator or co-transmitter because no agmatine-specific, postsynaptic receptor or agmatinergic system has been identified to date. The prevailing view is that the most common central nervous system disorders have a polygenic origin. Due to the widespread presence of agmatine in the peripheral and central nervous system, it is conjectured to be a “magical shotgun” - a non-selective drug with multiple targets, potentially leading to more effective treatments^11,12^. AGM function in the central nervous system includes antidepressant-like effects^13,14^, protection against schizophrenia^15^, improvement of cognitive dysfunction in Alzheimer’s disease^16^, and improvement of multiple sclerosis^17^.

However, the molecular mechanisms underlying agmatine function in the brain are unknown. This is primarily due to a lack of knowledge on the spatio-temporal effector function of agmatine *in vivo*. To address this gap, the development of a genetically encoded biosensor capable of visualizing agmatine localization and dynamics in a non-invasive manner is key for unravelling the role of agmatine in neuro(patho)physiology. Here, we present a fluorescence-based, genetically encoded AGM biosensor, consisting of an AGM binding-domain derived from the putrescine periplasmic binding-protein (PBP) PotF from *E. coli* fused to a super-folder circularly permuted GFP (sfcpGFP^18,19^). In the sensor the large conformational change of the PBP upon ligand binding is transmitted to the fluorescent protein by altering the chromophore environment and thus triggering a change in signal^20,21^. We chose PotF due to its promiscuous binding of biogenic amines characterized in prior studies^22, 23^. Our engineering employed a semi-rational approach combined with a medium throughput fluorescence screening to optimize linker positions and improve AGM specificity. We applied the final sensor AGMsen in different cell types *in vitro*, i.e., in primary neuronal cultures. Our results show that the sensor AGMsen can be used as the first of its kind to track agmatine in various experimental set-ups, which will allow to elucidate AGM function not only *in vitro* but also *in vivo*.

## Results

### Scaffold selection and initial sensor construction

Previously, we identified AGM as a natural ligand (K_D_ = 0.22 µM) for the periplasmic binding-protein PotF. The high affinity of PotF towards AGM can be explained by the perfect mimic of the binding mode of putrescine (PUT), as determined by crystallography (Figure 1a & b)^22^. To facilitate the engineering of high agmatine affinity and specificity by reengineering the binding pocket, we first established a sensor for the native ligand putrescine. This involved the incorporation of circular permuted green fluorescent protein (cpGFP, carrying mutation K12R) into the putrescine binding-protein PotF of *E. coli*. Entry sites were selected based on assumed major backbone displacement in the hinge region, as elucidated on the bound structure of *E. coli* PotF (1A99^24^, Figure S1). The majority of entry sites gave rise to sensors that responded to the addition of putrescine (PUT; Figure S2) while only the insertion at position 332 showed no response. While most of the sensors displayed a decrease in fluorescence upon PUT addition, only the sensor 328/332 exhibited a fluorescence increase of ΔF/F_0_ ∼0.3 upon the addition of 50 µM PUT. As we aimed for a sensor that increased its fluorescence upon binding, we proceeded with the sensor 328/332. Next, we explored the effect of different linkers between N-PotF and cpGFP as well as cpGFP and PotF-C. Changes were constrained to the two residues each exiting (N-linker) or entering PotF (C-linker). From this screen, A327L/E328I of the N-linker emerged as the most favorable variant while the C-linker remaining unaltered (N332/P333). Since the screen was based on measuring fluorescence changes after PUT addition to crude extract of *E. coli*, we introduced two further mutations (S87Y, F276W) as they reduce initial PUT affinity from the native nanomolar to the micromolar range^23^ to mitigate potential confounding effects of intracellular ligand. Next, we substituted the initially used cpGFP with the circular permuted variant of superfolder GFP^19^ (sfcpGFP). This preliminary sensor (PUTsen) then became the basis for engineering a sensor with high sensitivity and specificity for agmatine.

**Figure 1:**
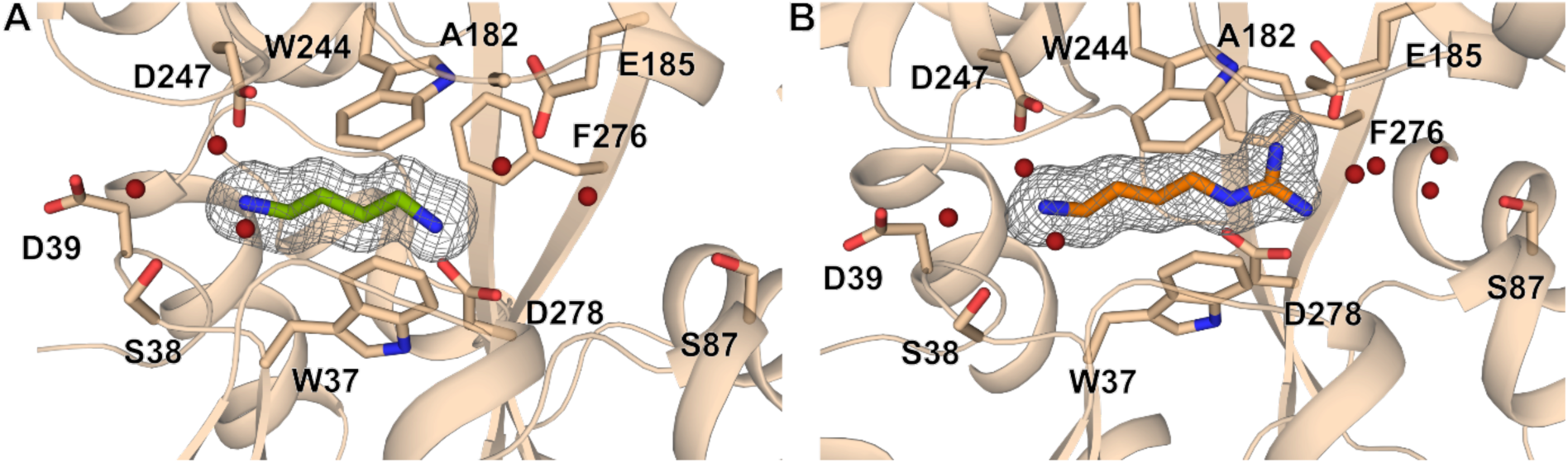
Overview of the binding pocket of PotF in complex with putrescine (A) and agmatine (B). Agmatine binds to wild-type PotF with high affinity as it mimics the putrescine binding mode. Ligand molecules and residues that form the binding pocket are shown as sticks. 2mFo-DFc maps for ligands are shown as gray mesh contoured at 1 σ using PyMOL.

### Tuning affinity and specificity of the sensor towards agmatine

In the next step, the PotF receptor module needs to be engineered for specific AGM binding because sensor performance will be assessed in cell culture experiments and possibly *in vivo* in the future. The natural biogenic amine ligands of wild-type PotF play an essential role in cell proliferation, will be present in experimental set-ups, and can even be included in media^25–28^. An unspecific sensor would thus recognize these other molecules, which can lead to high background levels or false positive results.

A high initial affinity to the target molecule is advantageous for engineering, since refining ligand specificity in a promiscuous binder like PotF often comes at the cost of losing affinity^23^. As a starting point for possible mutations, we first investigated PotF/D, a PotF variant into which the binding pocket of the homologous PotD was grafted^29^. To our surprise isothermal titration calorimetry (ITC) measurements revealed high AGM affinity in this variant (K_D_ = 4 µM, Figure S3 & Table S2). Additionally, PotF/D does no longer bind PUT and recognizes spermidine (SPD) with an affinity of 37 µM, making it a first construct with improved AGM specificity compared to PotF. Based on the thorough analysis of the differences between the binding pockets of PotF and PotF/D^23^, we randomly combined mutations (Table S1) identified in these studies. Subsequently, we started to screen biosensor constructs for affinity and specificity to AGM in *E. coli* lysate based on fluorescence changes upon ligand addition. The most interesting variants showed a substantial fluorescence increase in response to AGM coupled to low or no response to other PotF ligands such as PUT, SPD, and cadaverine (CDV). These variants were further purified by affinity chromatography to minimize the influence of lysate compounds and were re-screened for biogenic amine binding in a similar fashion. The outcome of this lysate and protein assay screening round, which led to the identification our final variant, is shown in Figure S4.

This final variant identified through our screening process carried the mutations S87Y and A182D. Additionally, we aimed to generate a non-binding control sensor to validate that changes in fluorescence are solely triggered by ligand binding. To achieve this, we introduced the mutation D247K strategically located in the first responding region of the protein to ligand encounter. By introducing a lysine at this position, we masked the primary amine binding-site in PotF generating a sensor that is incapable of binding any of the tested polyamines. We confirmed the effect of these mutations on AGM specificity by measuring binding affinities with ITC for the PotF receptor module without the fluorescence protein: PotF-S87Y-A182D displayed an affinity of 0.3 µM for AGM and no apparent K_D_ for the other tested ligands PUT, SPD, and CDV (Figure S3b). The control sensor showed no binding for any of the tested ligands (Figure S3c).

To examine and understand the binding of AGM further, we solved the crystal structure of PotF-S87Y-A182D in complex with AGM and compared it to the wild-type PotF structure. Interestingly, the two lobes of PotF are slightly more open in the S87Y-A182D construct (Figure 2a). We analyzed the relative orientation of the lobes in this structure as described before^22^ and found 10.9° wider opening and 7.1° wider twisting angles compared to wild type like closure. Still, most AGM interacting residues in the binding pocket do not differ (Figure 1b & 2b). Since the lobes are slightly further apart, distances between interacting residues and important binding pocket water molecules can change. These distances are compensated by the bulkier ligand AGM but are most probably detrimental for high affinity binding of slim ligands like PUT and SPD.

**Figure 2:**
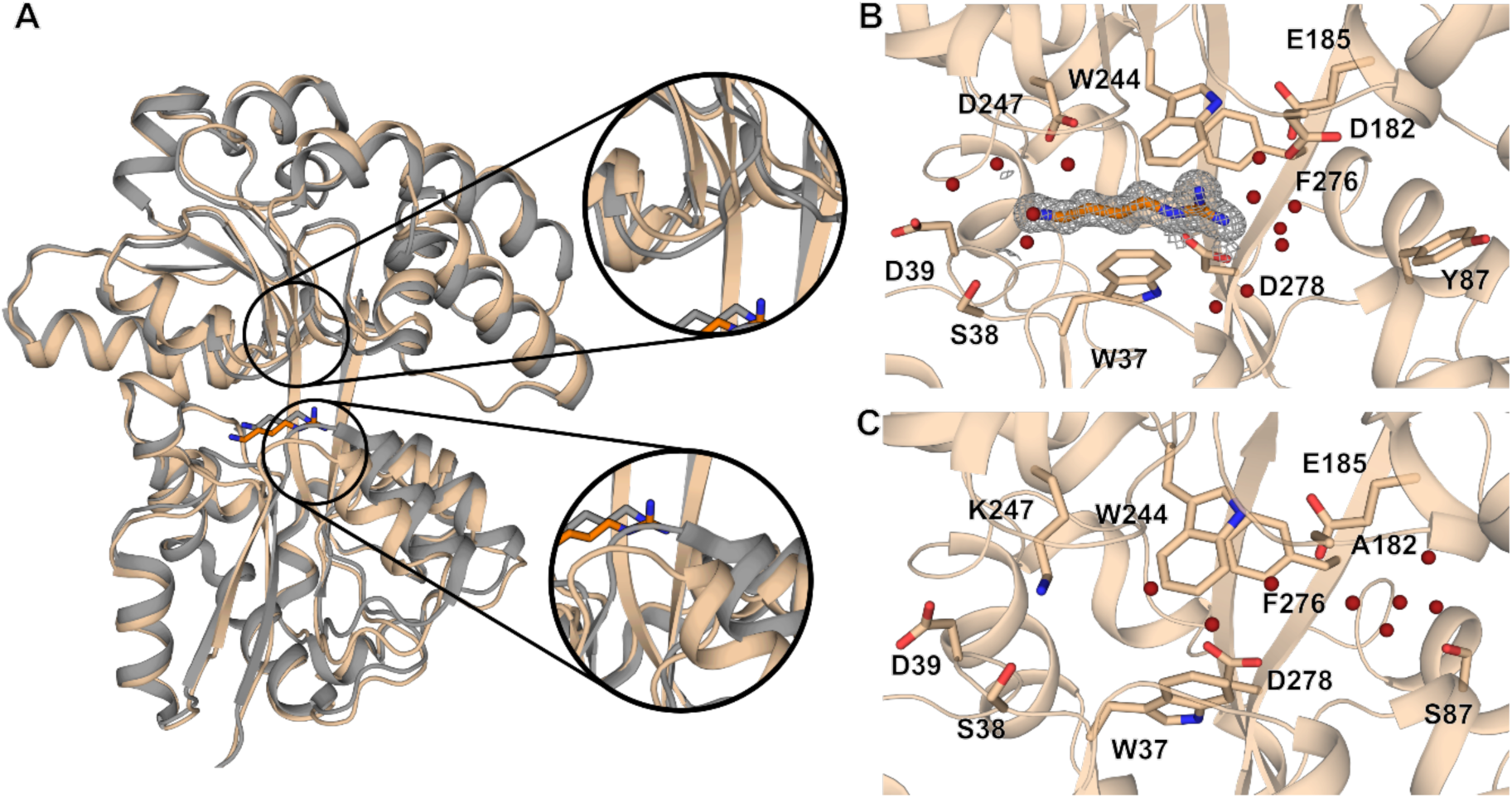
Alignment of the PotF receptor module (S87Y-A182D) of AGMsen in complex with AGM (pale orange) and PotF in complex with PUT (grey; A). Overview of the binding pocket of the PotF receptor module of AGMsen (B) and the control sensor (C). The structure of the AGMsen receptor module is slightly more open than wild type PotF. The positioning and interactions with agmatine in the pocket remained mainly unchanged compared to wild type PotF (Figure 1b). In the control sensor residue K247 occupies the primary amine binding site and, thereby, prevents ligand binding. Ligand molecules and residues forming the binding pocket are shown as sticks. 2mFo-DFc maps for ligands are shown as gray mesh contoured at 1 σ using PyMOL. Structure statistics can be found in table S4.

The control receptor module PotF-D247K showed no apparent binding to all tested polyamines in ITC (Figure S3c). Furthermore, the crystal structure confirmed that the amine of the newly introduced lysine 247 occupies the primary amine binding-site of the PotF pocket and, thereby, blocks ligand binding as expected (Figure 2c).

### Performance of the purified sensor

After the promising ITC results for the isolated receptor modules, we assessed the binding capabilities in the framework of the sensor with the added sfcpGFP. We named the resulting sensor containing S87Y and A182D AGMsen and with purified sensor conducted fluorescence-based dose-response measurements upon addition of AGM, PUT, SPD, and CDV at concentrations ranging from 0.1 µM to 10 mM. Here, AGMsen showed a dynamic range (ΔF_max_/F_0_) of 3.0 and an AGM affinity of about 38 µM (Figure 3a, Table S3). For the control sensor, only a small increase in fluorescence intensity at very high AGM concentrations was observable in the dose-response measurements, which corresponds to a K_D_ of 3.6 mM (Figure 3b, Table S3). Overall, PotF variants showed good performance in ITC and fluorescence-based dose-response measurements as sensors. The observed differences in affinity between the two techniques are likely due to the influence of the inserted GFP on the PotF receptor modules.

**Figure 3:**
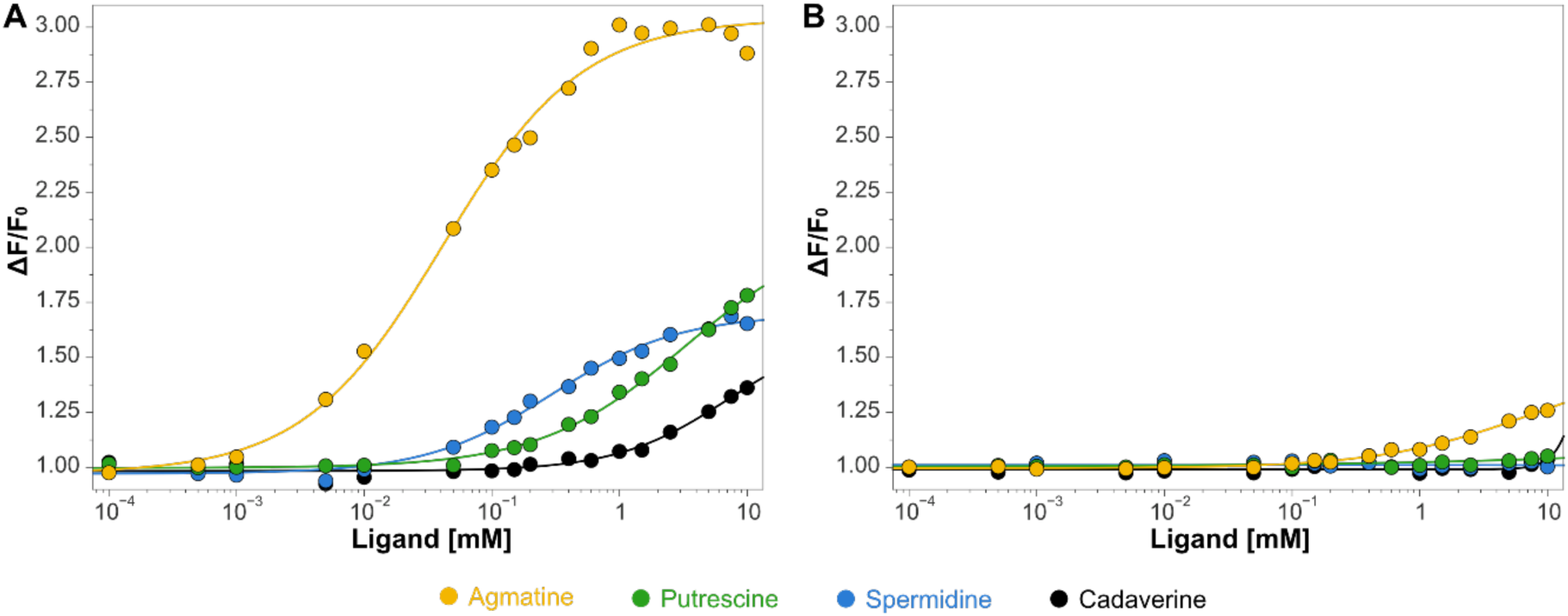
Dose-response curves in response to different biogenic amine ligands for purified AGMsen (A) and purified control sensor (B) from *E. coli*. AGMsen carrying mutations S87Y and A182D shows a better dynamic range of 3.0 and an affinity of 38 µM for AGM. Additionally, AGMsen shows low affinities for SPD, PUT and CDV with K_D_’s of 244 µM, 1.9 mM, and 3.4 mM, respectively. The control sensor (B) displays residual AGM affinity (3.6 mM) accompanied by a low dynamic range (ΔF_max_/F_0_ 1.3). Data points represent the mean of triplicates and the fit was done with the Hill equation using the fit-o-mat^30^. All K_D_ values can be found in Table S3.

### Sensor characterization in cells

We next introduced the sensors into the pcDNA3.1. and pDisplay vectors for eukaryotic expression to further characterize them in cell culture experiments. With pDisplay, we introduced an N-terminal signal peptide to guide the sensor to the secretory pathway and a C-terminal transmembrane domain to anchor the sensor. First, we performed immunocytochemistry to visualize the expression of AGMsen in the pcDNA3.1 and the pDisplay backbone in HEK293 cells. To analyze anchoring of the sensor to the outer cell membrane, we performed labeling with and without cell permeabilization. The expression of the sensor in both constructs was verified in permeabilized cells but membrane anchoring was only observed in the non-permeabilized cells expressing pDisplay-AGMsen (Figure 6a). Thus, the sensor is secreted and incorporated into the outer cell membrane.

To further characterize the sensor performance in different cell-based set-ups, we expressed AGMsen and the control sensor in HEK293 and performed dose-response measurements in cell lysate. The performance was then compared to the prior *E. coli* assays to ensure that the constructs behave similarly and are not influenced by specific eukaryotic features, e.g., posttranslational modifications. No notable differences for affinities of all ligands were observed (Figure 4b & c and Figure 3a &b, Table S3), thereby confirming our bacterial set-up as a robust screening platform that can be used to extrapolate sensor function in eukaryotic cells. The only observable difference is in the 20% lower maximum dynamic range upon AGM addition, which is expected as some lysate components might quench the fluorescent signal (Figure 4b).

**Figure 4:**
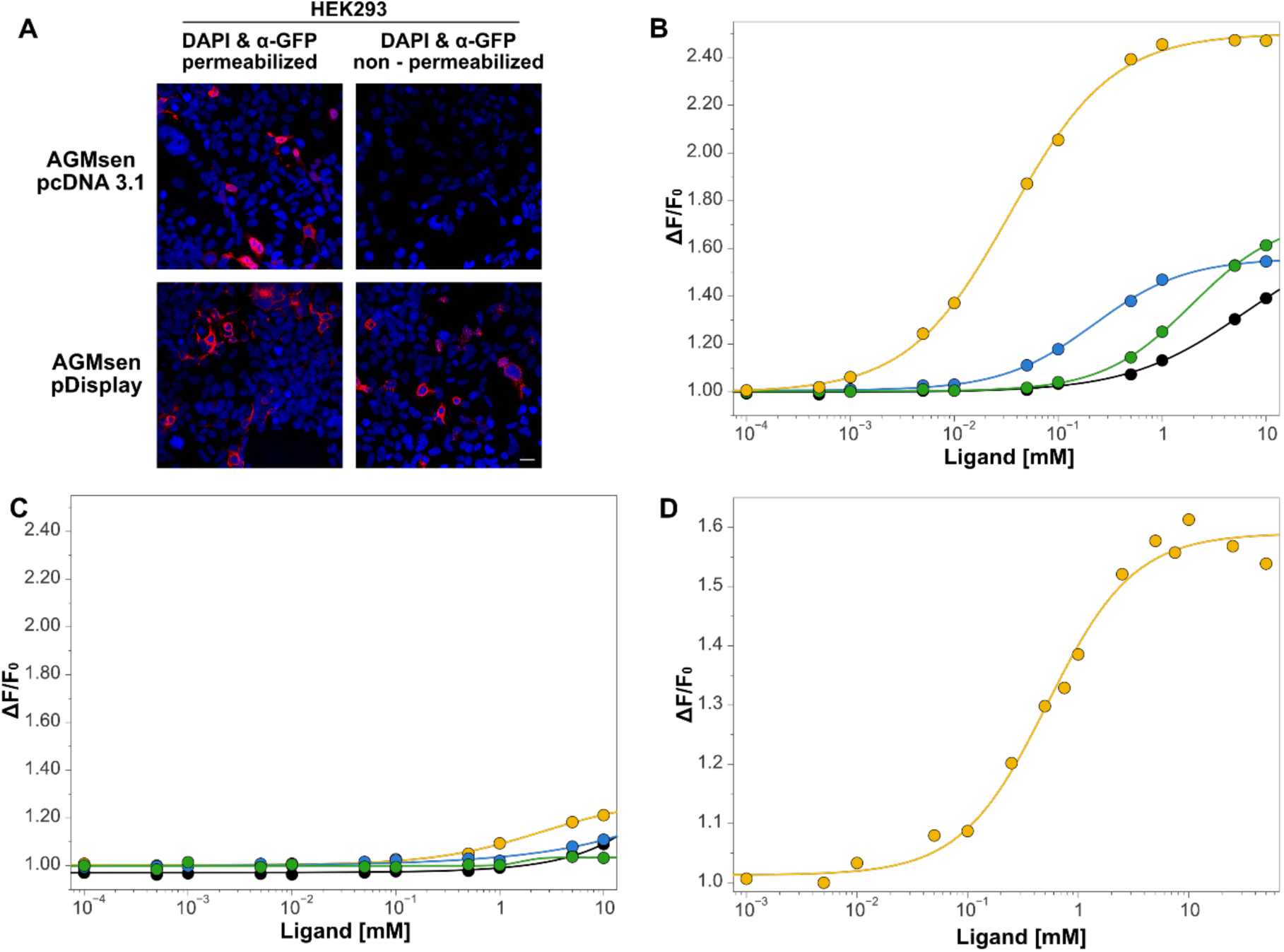
Immunocytochemistry of HEK293 cells expressing AGMsen in pcDNA3.1 or pDisplay, with or without permeabilization (A). Dose-response of AGMsen (B) and the control sensor (D) in HEK293 cell lysate. AGMsen was expressed in HEK293 cells using either the pcDNA3.1 or the pDisplay backbone (A). In permeabilized cells, GFP expression was detected in both cases, while in non-permeabilized cells, GFP was only observed for the pDisplay variant, confirming the correct incorporation of the sensor after secretion into the outer cell membrane. Cells are counterstained with DAPI. Scale bar = 20 µm. In HEK cell lysate the affinities for AGMsen are similar to measured affinities for purified sensor from *E. coli.* with the dynamic range being slightly lower (AGM K_D_= 35 µM, ΔF_max_/F_0_ = 2.4). The control sensor also performs the same displaying residual AGM affinity of 2.4 mM accompanied by a low dynamic range of 1.2. The response of AGMsen displayed on top of HEK293 cells (D) to AGM shows a reduced K_D_ (525 µM) and dynamic range (1.6) compared to the non-displayed variant. Data fits were done with the Hill equation using the fit-o-mat^30^. All K_D_ values can be found in Table S3.

Then, we measured dose-response curves for HEK293 live-cell suspensions in a PTI spectrofluorometer for pDisplay-AGMsen (Figure 4d). Here, the secreted sensor displayed a lower K_D_ of 525 µM and dynamic range of 1.6 compared to the lysate from intracellularly expressed sensor (K_D_ = 35 µM, ΔF_max_/F_0_ = 2.4, Figure 5c). Still, the sensors response is likely enough to visualize release events as neurotransmitters reach high concentrations in the synaptic cleft (as discussed below). Therefore, we subcloned pDisplay-AGMsen and the control sensor into a pAAV backbone with a synapsin promotor for neuronal expression and generated Adeno-associated virus containing the sensors. We next transduced primary rat hippocampal neurons with the pDisplay-AGMsen and control sensor and observed expression after 8 – 10 days (Figure S5). To quantify the AGM response, we analyzed neurons for each added AGM concentration using ImageJ as described in Figure S6. Results for different cells in the same experiment do scatter (Figure S7), nevertheless, clear trends are observable (Figure 5 & S7). Overall, fluorescence signal changes in neurons were much lower compared to HEK but the resulting K_D_ of about 700 µM is in the same regime as for the sensor displayed on HEK cells. This confirms functionality for AGMsen displayed on rat hippocampal neurons.

**Figure 5:**
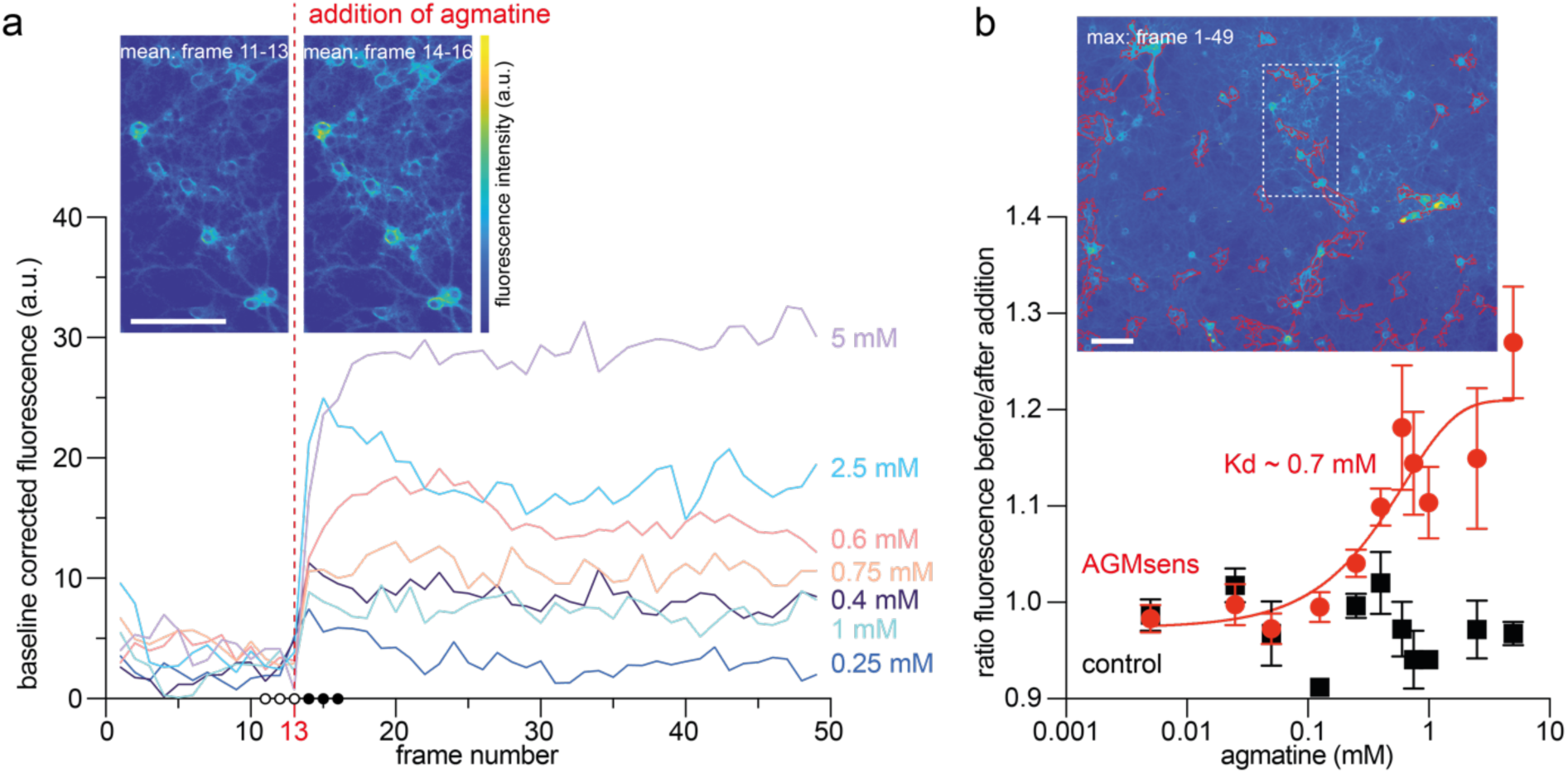
Response of AGMsen in neurons. AGMsen was expressed in rat hippocampal neurons and imaged using a Zeiss Observer.Z1 widefield microscope before and after the addition of agmatine (dashed red line in a). **A**: Exemplary curves of single ROIs after the addition of AGM at concentrations between 5 and 0.25 mM. Top left insets show magnifications of the 5 mM addition data (dashed white rectangle in **B**) with the mean of three frames before and after agmatine addition shown (hollow and filled circles on x-axis). **B:** Mean fluorescence change ratio of AGMsen and control sensor at different AGM concentrations. Shown is the mean pixel value of a minimum of 10 ROIs per datapoint. Error bars indicate the standard deviation. The AGMsen data can be fitted with a sigmoidal yielding a K_D_ of ∼0.7 mM (quality of fit r-squared 0.83). The inset shows a maximum intensity projection of the data for the 5 mM addition. The automatically chosen ROIs are indicated in red. All scale bars 100 µm. All colour bars as in inset **A**. Curves for AGMsen and control sensor for all concentrations and ROIs can be found in Figure S7 and a detailed description of the analysis procedure in Figure S6.

## Discussion

### Design aspects

In the engineering of the sensor the functional guanidino group proved to be a major factor to achieve selectivity for an otherwise indiscriminate polyamine-binding protein. The unconventional pairing of the high similarity in binding mode with the uniqueness of a different functional group was the key ingredient in the design process. In the crystal structure of the sensor, the positioning of AGM does not differ from the one in wild type suggesting that the other amines could still recognize the primary amine binding site with rudimentary affinity. The main structural changes occur in the distal site of the pocket (S87Y & A182D, Figure 1 & 2) where the guanidino group of AGM is located. It appears that this larger and bulkier group withstands the mutational changes and maintains high affinity while the smaller primary amine groups of other ligands lose most of their affinity. This discrepancy is likely caused by slightly incomplete closure of the PotF-S87Y-A182D receptor module in AGMsen, resulting in suboptimal coordination for less bulky ligands like PUT and SPD. It also has consequences for the sensor, whose functionality depends on the closing of PotF, and hence, might display a lower dynamic range than optimal closure would provide.

Interestingly, we observed discrepancies between the affinities of the actual sensor and the isolated receptor module determined by ITC. This might be attributed to different factors. On the one hand, the sensor is purified only by IMAC and mostly screened in lysate which can affect the performance of the receptor module. On the other hand, GFP with a size of 28 kDa is inserted into a region of PotF sensitive to movement and, therefore, can influence PotF by exerting forces or resistance on the closing mechanism. It is notable that AGMsen does not completely omit binding of the other amines, especially for the longest comparable ligand SPD. Still, fluorescence signal and affinity in purified sensor are much lower for SPD (245 µM), hence, this is not expected to influence the results since the sensor would always prefer AGM in an *in vivo* setting.

In addition, our knowledge of the PotF pocket and dynamics enabled us to construct a functional control sensor. The D247K mutation occupies the primary amine binding site, where typically all characterized ligands of PotF bind in the same fashion. The crystal structure displays a nearly fully closed conformation which could explain the good signal of the control sensor. This allows for reliable controls and the assessment of robustness for future experimental set-ups since drifts or other issues in fluorescence signal will be easily traceable with the control sensor.

### Sensor performance

Comparisons of the assay results from *E. coli* and HEK cells (Figure 3 & 4) confirm the robustness of our screening system and alleviate concerns regarding potential impact of codon usage on the results. Interestingly, the HEK displayed AGMsen showed a lower apparent affinity and dynamic range than the non-displayed construct, suggesting that the dynamics of the sensor are influenced by the addition of the secretion signal, linkers, and transmembrane domain. This effect is not unexpected as these modifications, similar to the insertion of sfcpGFP, can also influence the dynamics and closing mechanism of PotF. Based on our ITC measurements that indicate a potential K_D_ of 0.3 µM for the AGMsen receptor variant, further optimization of the sensor might be achieved by changing the different linkers to obtain better signals and provide less influence on the affinity, especially at lower ligand concentrations. Moreover, the epitope tags from the construct could be removed as in pMinDisplay, a version of pDisplay missing the HA tag^4^. Another factor that is important to address are reaction conditions: the displayed sensor ideally works in an extracellular environment whereas the non-displayed sensor functions intracellularly, meaning a different experimental set-up is required i.e., changing salt content and concentrations. Given that ligand binding in the PotF pocket is highly mediated by coulomb forces between ligand amines and carboxyl harboring sidechains^22^, variations in ionic strength of the buffer could potentially impact affinities. However, it is noteworthy that the highest affinity (0.3 µM, Figure 3, Table S2) for the PotF receptor module was determined by ITC in a high salt buffer containing 300 mM NaCl, hence we believe that the influence of the different extensions on PotF plays a bigger role than slight differences in buffer conditions.

The importance of improving sensor dynamics further becomes apparent when assessing the neuron data. Due to generally lower signal increase, background noise can have a stronger influence on the results and neuron culture tend to be more inconsistent overall. This could be counteracted by a stronger, more prominent signal. The response of AGMsen displayed on neurons was slightly lower but in a similar regime as observed for the HEK displayed AGMsen. One reason could be the difficulties with signal intensity and high fluctuations between cells in the same measurements, resulting in elevated errors in neuron culture. Additionally, the use of neurons from rats, a different organism, could also lead to small changes in performance. Nevertheless, the HEK cell and rat neuron displayed AGMsen are comparable suggesting that newly optimized constructs could be easily characterized and compared in HEK cells first before moving to other organisms and experimental set-ups to save time and resources.

### Conclusion

Leveraging our understanding of PotF, an AGM sensor was successfully engineered that functions effectively in both HEK cells and rat hippocampal neurons. This sensor is available in multiple vectors for expression inside cells (pcDNA 3.1) or for display on cells (pDisplay and pAAV-pDisplay). To directly confirm the authenticity of signal changes in AGMsen being induced by AGM binding and no other components of the experimental set-up, we also provide a non-binding control sensor. While there is room for improvement regarding dynamic range and affinities in the current sensor to track weaker AGM signals, these are minor imperfections that can be addressed through engineering in the future. For AGM to fulfill its neuromodulatory functions, like other transmitters, high concentrations in the synaptic cleft are needed. It has been shown that AGM is colocalized with glutamate, the major excitatory neurotransmitter of the CNS, in the hippocampus^31^. The concentration of glutamate at cultured hippocampal synapses peaks at 1.1 mM^32^ and it has been described that concentrations surpass 1 mM after stimulation by an action potential^33^. It is anticipated that AGM is reaching concentrations in this range which is reliably detectable by our sensor. Moreover, the high µM affinity ensures that the sensor is not affected by small fluctuations in AGM concentrations and, hence, will only signal genuine release events. Therefore, AGMsen represents an important first step toward better understanding AGM-specific mechanisms in a non-invasive manner and provided a useful tool through close collaboration with users can be tuned further towards diverse research needs.

## Methods

### Initial construction in pET21b(+)

Initial sensor construction was conducted by a two-step splicing overlap extension PCR with Phusion polymerase (New England Biolabs) using standard protocol with an adjusted elongation time of 50 s. Used oligonucleotides can be found in Table S5. Resulting fragments were purified after agarose gel electrophoresis. After subsequent restriction digest with *Nde*I and *Xho*I (both from Thermo Fisher Scientific) the fragment was ligated with equivalently linearized pET21b(+). Top10 cells were transformed with the reaction mixture via heat shock. DNA of overnight cultures was isolated using NucleoSpin® Plasmid EasyPure-Kit (Machery & Nagel) according to the manufacturers protocol and implementation of mutations was confirmed by sequencing.

### Targeted mutagenesis

Mutations for the construction of different AGMsen and PotF variants were introduced by a modified QuickChange PCR using KAPA® polymerase (Roche) followed by an additional ligation step with T4 DNA ligase (New England Biolabs). All used oligonucleotides for mutations presented in this work are listed in table S5. Transformation, DNA isolation and construct verification was conducted as described above.

### Cloning into pDisplay

Sensor constructs were amplified by a standard Q5® High-fidelity DNA-Polymerase protocol (New England Biolabs) using oligonucleotides (Table S6) that introduced *Sfl*I and *Pst*I (both from Thermo Fisher Scientific) restriction sites. After restriction using standard protocol for the respective enzymes and subsequent purification, fragments were ligated with equivalently linearized pDisplay vector. Transformation, DNA isolation and construct verification was conducted as described above.

### Cloning into pcDNA

Final and control sensor fragments were cloned into pcDNA using Gibson assembly. Fragments were amplified using a standard Q5® High-fidelity DNA-Polymerase protocol (New England Biolabs). Primers were designed using NEBuilder® and are shown in table S6. The PCR added a Kozak sequence to facilitate translation in eukaryotic cells. Amplified vector and insert fragments were purified from 1% (w/v) agarose gels using NucleoSpin® Gel and PCR Clean-up kit (Macherey & Nagel) according to manufacturer’s protocol. Subsequently, fragments were mixed in a 1:1 ratio (n/n) to a final DNA quantity of 200 ng in 5µl reaction volume. DNA mix was added to Gibson Assembly reaction mix and incubated for 1 h at 50°C. Transformation, DNA isolation and verification was done as described above.

### Cloning of sensor genes into an adeno associated virus (AAV) vector

For expression on the surface of neurons, the genes for the displayed AGMsen and the control sensor were placed under the control of the human synapsin I promoter. Therefore, both sensors were excised from the respective pDisplay-plasmids including the N-terminal Igk leader and HA, as well as C-terminal Myc and PDGFR domain sequences using *Xba*I and *Hind*III. Afterwards the genes were subcloned into an AAV vector (rAAV-Syn1-MCS; serotype 2/1) and used for recombinant AAV production.

### Protein expression and purification

Receptor modules were expressed and purified as described in Kröger et al 2021a &b.^22,23^ Sensor variants were overexpressed in autoinduction medium ZYM-5052^34^ for 18 h at 30°C and 180 rpm using *E. coli* BL21 (DE3) cells. After expression, cultures were pelleted (3500 g, 20 min, 4°C) and resuspended in 20 mL buffer S1 (20 mM Tris pH 8, 100 mM NaCl, 20 mM imidazole) per pellet of 500 mL culture. Cells were disrupted by sonication (Branson medium tip, 2 x 3 min, Duty 40%, Output: 4). After centrifugation (40.000 g, 1 h, 4°C) supernatant containing C-terminally His_6_-tagged target protein was loaded onto a 1 mL HisTrap™ FF column equilibrated with 10 CV buffer S1. The column was washed with 10 CV buffer S1 followed by elution with buffer S2 (20 mM Tris pH 8, 100 mM NaCl, 500 mM Imidazole). Elution of the sensor was tracked by the greenish hue. Afterwards, buffer was exchanged to desired assay buffer using NAP 25 columns. Purified sensor was stored at 4°C if not used immediately.

### Isothermal Titration Calorimetry (ITC) of receptor modules

ITC was performed as described in Kröger et al 2021a & b^22,23^. Used protein and ligand concentrations can be found in table S7.

### Crystallography of receptor modules

All crystallization experiments were set up as sitting drop vapour diffusion experiments in 3-well Intelli plates (Art Robbins Instruments) using a protein concentration of 40 mg/mL (PotF-S87Y-A182D) and 30 mg/mL PotF-D247K and a 20-fold molar excess of ligand if added. Protein-ligand mixtures were equilibrated at 293 K for several hours before crystallization setups.

Crystals for PotF-S87Y-A182D were obtained in 0.085 M sodium acetate pH 4.6, 0.17M ammonium acetate, 30% PEG 4000 and 15% glycerol and for PotF-D247K in 2.4 M ammonium sulfate, 0.1 M Bicine pH 9.0 and 10% Jeffamine M600.

Crystals were mounted using CryoLoops and then flash-cooled in liquid nitrogen. In case of PotF-D247K crystals were first transferred into a cryogenic solution made of reservoir solution and 1.7 M malonate matching the pH of the condition. Data collection at 100 K was done at beamline BL 14.1 at the synchrotron BESSY II, Helmholtz-Zentrum Berlin^35^. Diffraction data was processed as described in Kröger et al 2021a & b^22,23^.

### Lysate-based fluorescence assay

Sensor variants were overexpressed in 10 mL autoinduction medium for 18 h at 30°C and 180 rpm. Afterwards, bacteria were pelleted by centrifugation (20 min, 3500 g, 4°C) and resuspended in 2 mL of assay buffer (20 mM Tris pH8, 100 mM NaCl). The suspension was transferred to two 1.2 mL tubes of an eight’s-cluster-strip (biolab products) and 180 - 220 µg of 0.1 mm glass beads (biolab products) were added. Cells were disrupted by using a bead mill (Bead ruptor, Omni®) 24 at 6 m/s for ten cycles of 30 s with 1 min breaks between each cycle. Debris and beads were pelleted by centrifugation (20 min, 4800 g, 4°C). 180 µL of supernatant was transferred to a Nunclon F96 MicroWell™ black plate and a first fluorescence intensity measurement (λ_ex_ = 470/10 nm, λ_em_ = 515/10 nm) was conducted to adjust the gain for the tested constructs to a fluorescence of roughly 10000 - 20000 units. The detection method of the reader (Tecan Spark) was set to fluorescence top reading, the gain according to the initial scan, the excitation and emission as stated before, and the Z-Position to 16800 µm. Solutions of different ligands were prepared in assay buffer. For each well the initial fluorescence intensity was measured, followed by 10 x 5 µL injections of 100 mM ligand solution using injector pumps. After each injection, a 2 mm and 150 rpm double orbital shake was conducted for 5 s prior to the measurement. As a control, buffer was added in an additional well in 10 x 5 µL injections. The measured values were corrected by the fold-change in fluorescence intensity of the buffer control and the resulting fold increase in fluorescence intensity was plotted against the ligand concentration to evaluate ligand binding.

### Protein-based fluorescence assay

Plate reader setting and data evaluation were kept consistent with the lysate assay. First, a dilution series (1:10 - 1:10.000) of the sensor was measured in the respective assay buffer (20 mM Tris pH8, 100 mM NaCl) in a total reaction volume of 180 µL to adjust fluorescence emission to 10000 –20000 fluorescence counts. Fluorescence intensity was measured as triplicates. For each well the initial fluorescence intensity was measured, followed by 5 x 5 µL injections of 20 mM ligand solution and 9 x 5 µL injections of 100 mM ligand solution using injector pumps. After each injection, a 2 mm and 150 rpm double orbital shake was conducted for 5 s prior to the measurement. As a control, buffer was added in an additional well in 14 x 5 µL injections.

### Dose-response assay in E. coli

Plate reader settings and dilutions were kept consistent with the other fluorescence assay. To measure binding affinities of the final sensor variant, 180 µl of adequate sensor dilution was combined with 20 µl of ligand solutions ranging from 0.01 mM to 500 mM. Reaction mix was incubated for 5 - 10 min at RT to ensure endpoint measurements. A buffer was chosen that represents HEK cell lysate assay buffer (10 mM NaCl, 5 mM KCl, 2 mM MgCl_2_*6H_2_O, 2 mM CaCl_2_*2H_2_O, 10 mM glucose &10 mM HEPES pH 7.4). Data was normalized to the buffer measurement and fluorescence increase was plotted against ligand concentration. Data was fitted sigmoidal for a 1:1 binding ratio of sensor and ligand using the Hill-equation.

### Preparation of primary hippocampal cultures

Primary hippocampal neurons were cultured from C57BL/6 mice as described before^36^. Hippocampi were dissected from mice at embryonic day 15-19 (animal license protocol AZ 81-02.04.2020.A100, LANUV, NRW, Germany), washed with HBSS (Life Technologies), and incubated with 0.025 g/ml trypsin (Life Technologies) for 20 min at 37°C. After washing with HBSS, DNA was digested using 0.001 g/ml DNase I (Roche) and the tissue was further dissociated using filter tips. The dissociated cells were seeded on coverslips coated with Poly-D-lysine (Sigma-Aldrich) in a 24-well plate at a density of 40.000 cells per well. The cells were incubated in Neurobasal Medium supplemented with 2% B-27 and 1 mM L-glutamine or in Basal Medium Eagle supplemented with 2% B-27, 0.5% glucose, 0.5 mM L-glutamine, and 1% fetal bovine serum (ThermoFisher Scientific) at 37°C with 5% CO_2_ until further use.

### Recombinant adeno-associated virus (rAAV) production

rAAV of serotype 2/1 were produced as described before^37^. Briefly, HEK293T cells were transfected using the calcium-phosphate method with pAAV-Syn-Display-AGMsen and pAAV-Syn-Display-control sensor together with the adenoviral helper plasmid pFΔ6, pRV1, and pH21, the latter encoding rep and cap genes. Cells were harvested 48 hours after transfection and lysed in 0.5% sodium deoxycholate (Sigma-Aldrich) and 50 units/ml Benzonase endonuclease (Millipore). Viral particles were purified by HiTrap^TM^ heparin columns (GE Healthcare) and concentrated with Amicon Ultra Centrifugal Filters (Millipore) to a final volume of 400 µl. An aliquot of the virus was validated on an SDS-polyacrylamide gel by Coomassie Blue staining.

### Transfection of HEK cells and transduction of hippocampal neurons

HEK293 (CRL-1573) cells were obtained from American Type Culture Collection (ATCC) and grown in Dulbecco’s Modified Eagle Medium (DMEM, Gibco) supplemented with 10 % FCS to 90 - 95% confluency. Cells were transfected with 9.5 µg sensor plasmid per 9 cm petri dish and 0.5 µg per 4 well plate. Therefore, DNA and polyethyleneimine (PEI, Sigma Aldrich) were mixed in OptiMEM (Gibco) to a final concentration of 10 ng/µl and 20 ng/µl, respectively. Reaction mix was incubated for 10 min at RT before adding it to the culture. Prior to that, old media was removed from the culture and replaced with 4x the volume to the reaction mix of fresh medium containing only 2% serum. Cells were left for 2 days at 37°C and 5% CO_2_ before further experiments to ensure good sensor expression. For transduction 0.5 - 1 µl of AAV was added directly into BME medium 4 days after isolation and incubated for 8 - 10 days at 37°C and 5% CO_2_.

### Dose-response assay in HEK lysate

For harvest media was removed, and cells were washed with 1x PBS. Afterwards 1 ml of 1xPBS was added to the dish, cells were scraped and transferred to 1.5 ml tube. After centrifugation (500g, 5 min, RT) PBS was removed and replaced with assay buffer (10 mM NaCl, 5 mM KCl, 2 mM MgCl_2_*6H_2_O, 2 mM CaCl_2_*2H_2_O, 10 mM glucose & 10 mM HEPES pH 7.4) for hypotonic lysis. Cells were incubated for 15 min on ice and sonicated in a water bath for 1x 30 s and 2x 10 s to assist lysis. Afterwards cells were incubated on ice for 5 min, followed by vortexing thoroughly and another 5 min incubation on ice. Finally, debris was pelleted by centrifugation (1000 g, 10 min, 4°C) and supernatant was diluted 1:10 for the lysate assay. 180 µl of lysate was transferred to black walled fluorescence plates (Grainer) and 20 µl ligand solutions with concentrations ranging from 0.001 mM to 100 mM were added. Mixture was incubated for 5 min to ensure endpoint measurements. Plate was transferred to a FluoStar Omega plate reader (BMG Labtech) and whole well fluorescence was determined as follows: λ_ex_ = 485, λ_em_ = 520, Flashes 20, top optics, Gain set to 90% fluorescence in the well with highest AGM concentration. Data was normalized as described before for the *E. coli* assay.

### Dose-response assay for HEK displayed sensor

Fist, media was removed, and cells were washed with 1xPBS followed by incubation with 1 ml 1xPBS/EDTA for 15 - 30 min (37°C, 5%CO_2_) or until cells start to detach. Cells were flushed from the plate and spun down (500g, 5 min, RT). After discarding the PBS/EDTA solution, cells were resuspended in 1x HBSS (Gibco). Cells were diluted 1:20 and 1.8 ml of the solution was transferred to a 2 ml reaction tube to which either 200 µl buffer or ligand solution ranging from 0.01 mM to 500 mM were added. The mixture was incubated at RT for 5 min to ensure endpoint measurements. Afterwards it was transferred to a fluorescence cuvette. Each cell solution was measured in a PTI QuantaMaster spectrofluorometer (Horiba) at 20°C and λ_ex_ = 488/7.5 nm, λ_em_ = 512/5 nm for 5 seconds under constant stirring to ensure homogeneity during the measurement. For each concentration the measurements over 5 s were averaged and data was normalized as described before.

### Immunocytochemistry with and without permeabilization

For staining, HEK cells and hippocampal neurons were grown on coverslips in four well or 24 well plates, respectively. Without permeabilization primary antibody (ab6556 Anti-GFP, Abcam, 1:500) was added to the medium (DMEM, 10% FCS) and cells were incubated for 30 min at 37°C and 5% CO_2_. Afterwards cells were washed with PBS followed by fixation with 4% paraformaldehyde (PFA, Alfa Aesar) and another wash. Staining was blocked for 30 min with CT buffer (0.5% Triton X-100, 5% ChemiBLOCKER in 0.1M NaP pH 7, Sigma Aldrich & Merck) at RT. Secondary antibody (Goat anti-Rabbit A647 IgG, A21245 Invitrogen 1:500 or 111-604-144 Jackson Immuno 1:400) was added in CT buffer simultaneously with DAPI (4’, 6-Diamidino-2-Phenyl-indole dihydrochloride, 1:10000) as a DNA counterstain and cells were incubated for 45 min in the dark. Cells were washed with PBS, and coverslips were placed face down on microscope slides with a drop Aqua-Poly/mount (Tebu-Bio) mounting medium being left to harden for 24 h at 4°C. For staining with permeabilization the primary antibody is added after fixation and blocking with an additional PBS washing step between the antibodies. Confocal images were recorded with a Leica Sp5 microscope using the 63x magnification HCX PL APO lambda blue objective. DAPI and AF647 were imaged at 30% laser power with a 488 nm argon laser and a 633 nm HeNe laser, respectively.

### Live cell imaging of neurons expressing the sensor in response to Agmatine

After transduction and expression, the activity of the sensor was analysed using a Zeiss Observer.Z1 widefield microscope equipped with an Axiocam 506 trans imaging device. GFP fluorescence (λ_ex_: 450- 490 nm, λ_em_: 500-550 nm) was imaged using the 10x magnification LD EC PlanNeoFluor objective. For the measurements, medium was replaced with 150 µl ES buffer, followed by recording a GFP baseline for 2 min in 10 s intervals. Different concentrations of agmatine were added to final concentrations ranging from 5 µM to 5 mM in a 1:2 dilution and measurements were continued further for 6 min.

### Data Availability

X-ray coordinates of all solved structures have been deposited at the protein databank with accession codes: 8ASZ, 8AT0

## Acknowledgment

We thank Susanne Gillig and Simon Prisner for exploring early sensor constructs and Cornelius Fischer for help in testing agmatine sensor constructs. We acknowledge financial support and allocation of synchrotron beamtime by HZB and thank the beamline staff at BESSY for support. rAAV-Syn1-MCS was kindly provided by Martin Schwarz (Bonn University Medical School). All imaging was performed at the Microscopy facility of the University Hospital Bonn. This work was supported by the Deutsche Forschungsgemeinschaft grant HO4022/2-3, the Max-Planck Society and Universität Bayreuth.

## Supplementary Information

**Figure S1:**
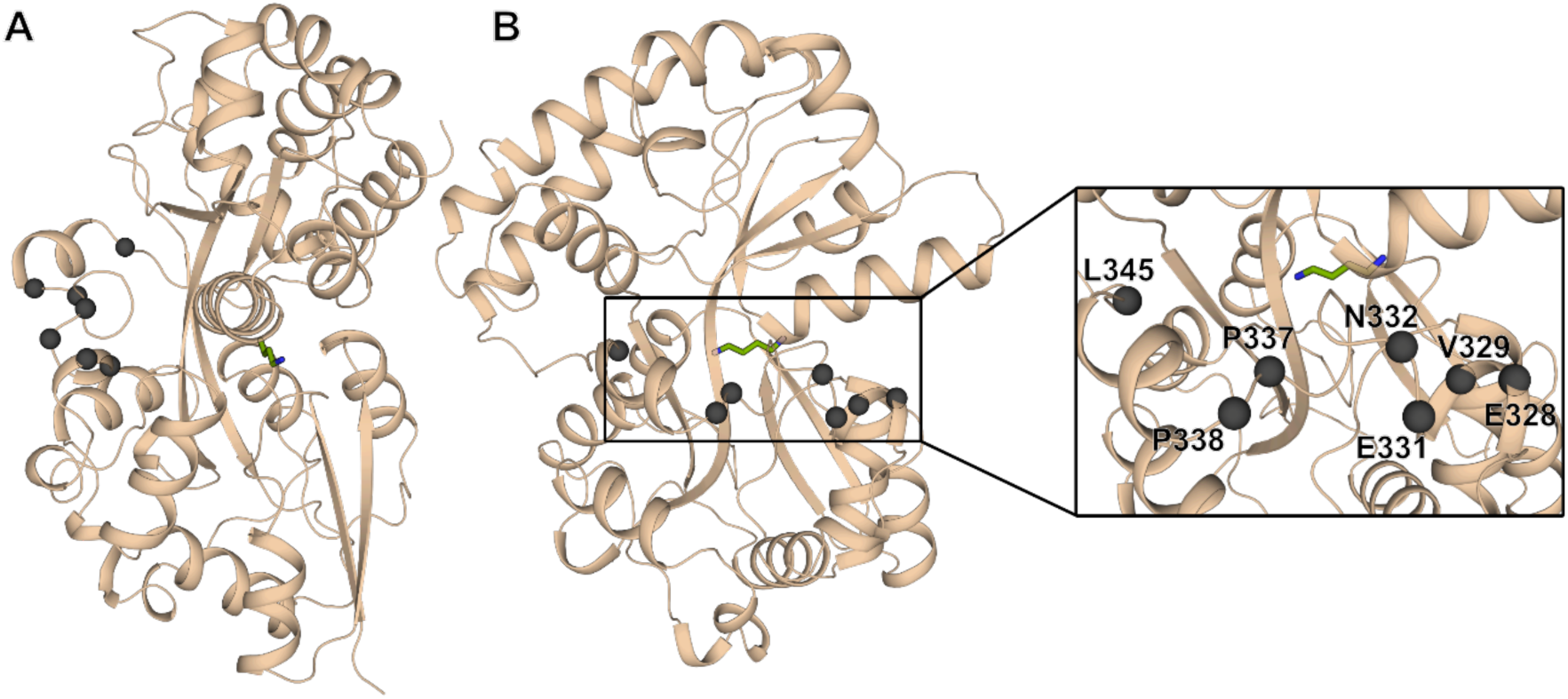
Insertion sites of cpGFP in PotF. Side (A) and back view (B) of PotF highlighting the insertion sites for cpGFP, which are located on the back side of the binding pocket close to the hinge region to create a motion sensitive sensor similar to other PBP-based sensors^20^. The Cα of the residues that were targeted for the insertion are shown as dark grey spheres. Putrescine is shown as green sticks to indicate the position of the binding pocket in relation to the cpGFP insertion region.

**Figure S2:**
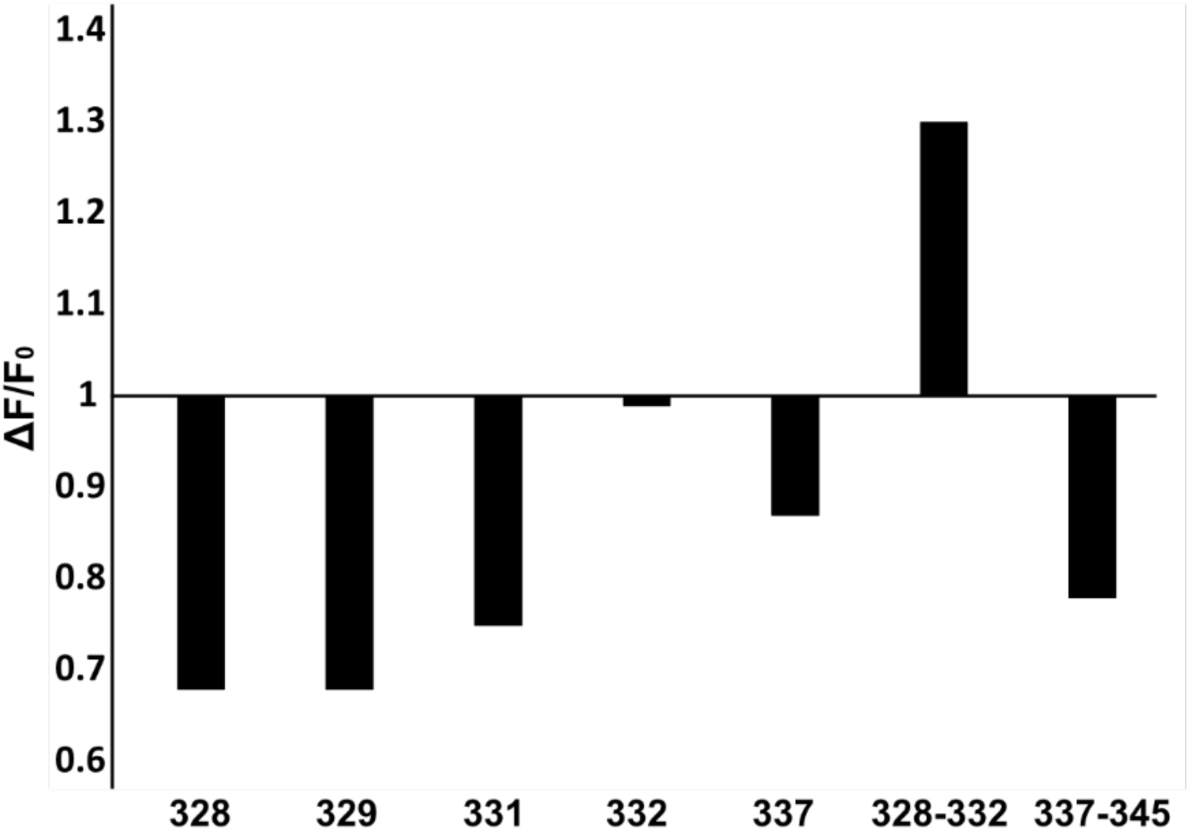
Fluorescence signal of the sensor constructs named after the cpGFP insertion sites upon putrescine addition. All single insertion sites result in a negative fluorescence signal, with position 332 not responding in general. Only the insertion in between positions 328 and 332 resulted in the desired gain in fluorescence upon adding 50 – 100 µM of putrescine.

**Figure S3:**
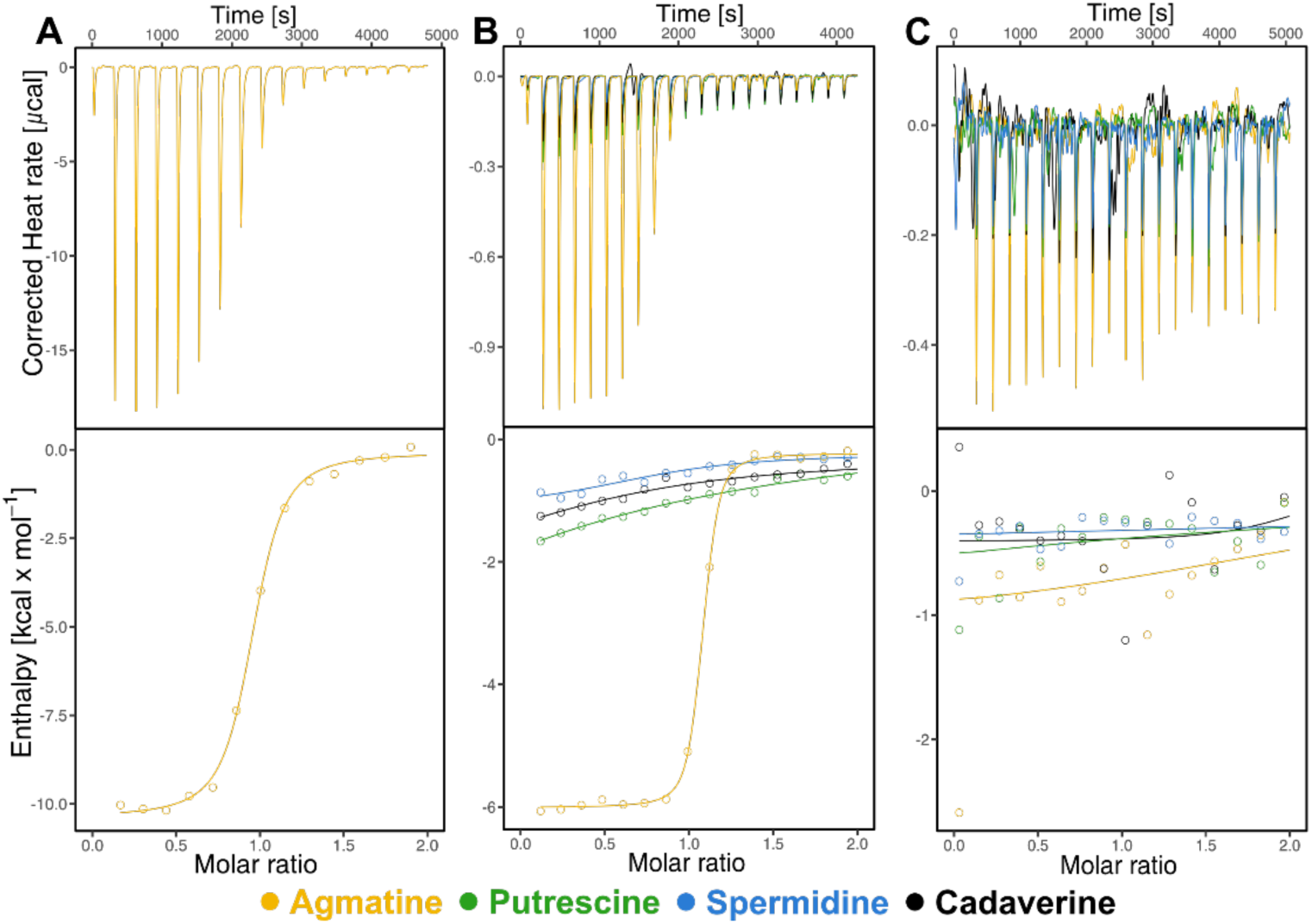
ITC measurements for PotF/D, PotF-S87Y-A182D and PotF-D247K. (A) PotF/D displays a K_D_ of 4 µM for AGM. (B) The PotF-S87Y-A182D receptor molecules display an affinity of 0.3 µM. Any binding of other polyamines was not detectable in ITC. (C) The designed control receptor PotF-D247K shows no binding of any tested ligand. Thermodynamics and stoichiometry of all measurements can be found in Table S2.

**Figure S4:**
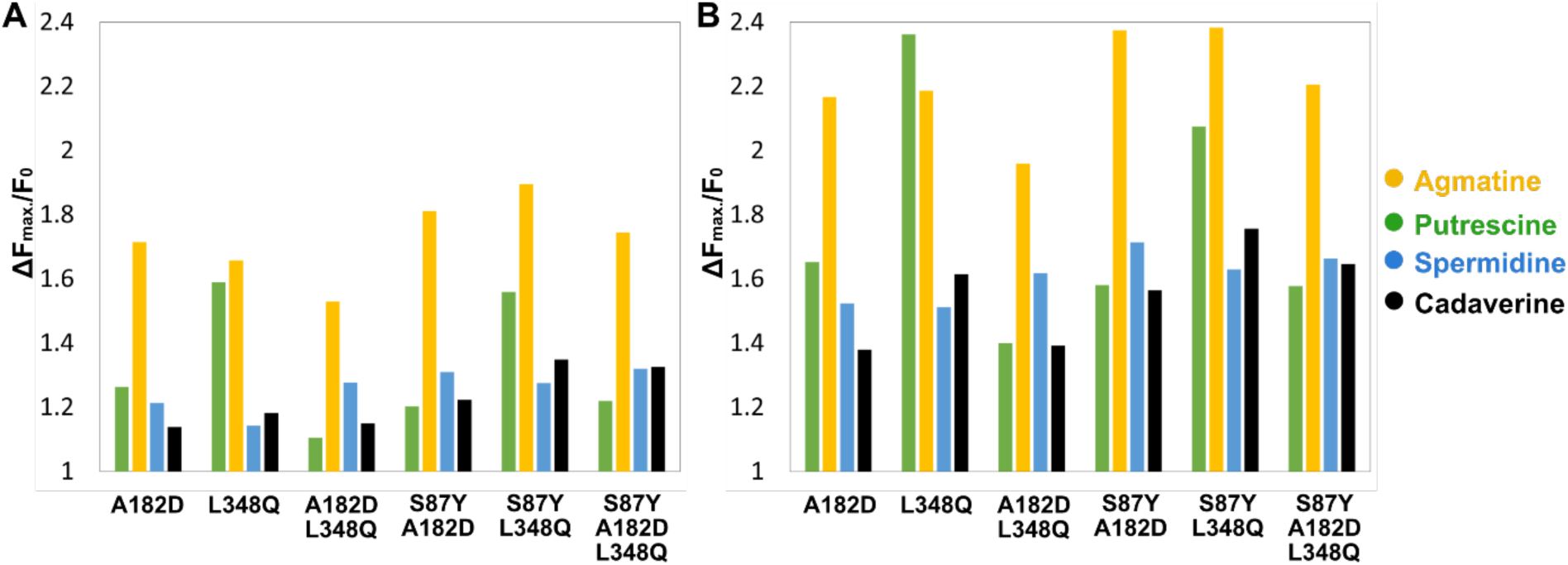
Lysate and purified protein screening assays. Examples of lysate (A) and purified protein (B) assays are shown for the same AGM sensor constructs. Overall, the fluorescence signal is similar albeit lower in the lysate screening. This representation only shows the maximum fluorescence gain after the last ligand addition. Protein assays are conducted to confirm lysate screening results and to lower the influence of possible endogenous biogenic amines from *E. coli* and other lysate components. S87Y-A182D is the variant that ultimately became AGMsen.

**Figure S5:**
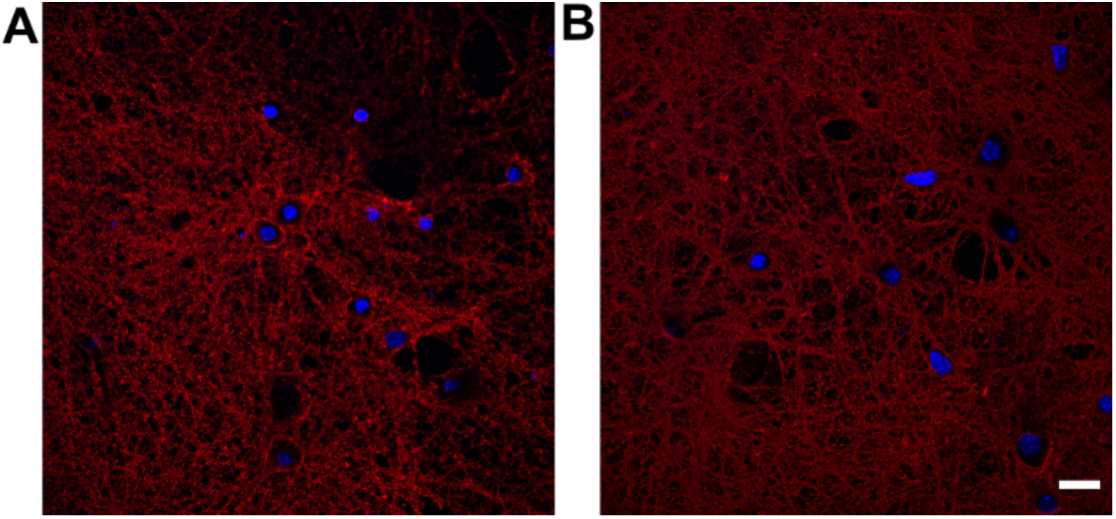
Antibody staining of primary rat hippocampal neurons. Neurons were transduced with pAAV-Syn-Display- AGMsen (A) or the control sensor (B). Both sensors express in neurons and are located in the outer membrane as confirmed by α-GFP antibody staining without permeabilization. Nucleus is counterstained with DAPI. Scale bar corresponds to 20 µm.

**Figure S6:**
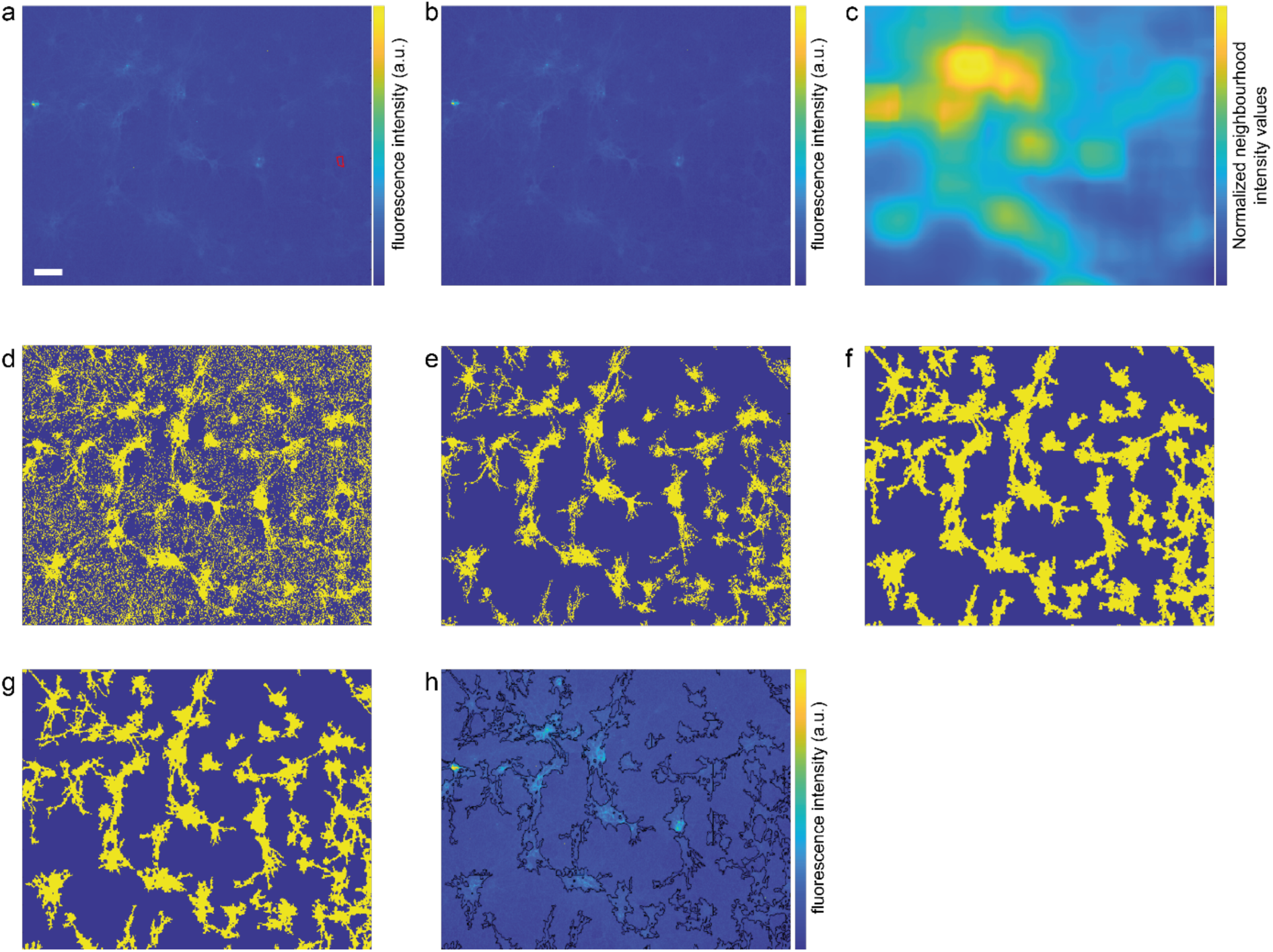
Analysis routine for neuronal imaging. All images shown are exemplary outputs of the described steps. Based on a maximum intensity projection of the tiff stack (**A**) a region with apparent homogenous minimal signal was chosen manually (red ROI). For each frame the mean pixel values of the background ROI were calculated and subtracted yielding a background corrected tiff-stack. On a maximum intensity projection of this stack an adaptive threshold algorithm was used (Matlab R2022a, “adaptthresh”, threshold setting 0.5, if not indicated differently all functions were used with settings as per default of R2022a) yielding a normalized map of neighborhood intensities (**C**). This map was used to binarize the maximum intensity projection of the background corrected stack yielding initial ROIs for the neuronal cells (“imbinarize”, **D**). Subsequently, small regions below 5000 pixels were removed (“bwareaopen”, **E**), the remaining ROIs were dilated (“dilate”, **F**) and subsequently eroded (“erode”, **G**) with a disk as morphological structuring element with a radius of 4 pixels (approximated with 6 lines, “strel”) yielding the final ROIs (**G**). In **H** the final ROIs are shown overlaying the maximum intensity projection of the background corrected tiff-stack. All images with scalebars as in **A** of 100 µm. The ROIs were used individually on each tiff-stack to extract the mean pixel values. Concatenation of mean pixel values yielded the time-trace for a given ROI. For each experimental condition 10-20 ROIs were analyzed. For each trace the change in fluorescence was calculated by dividing the mean of three frames after addition (14-16) by the three frames before addition (11-13).

**Figure S7:**
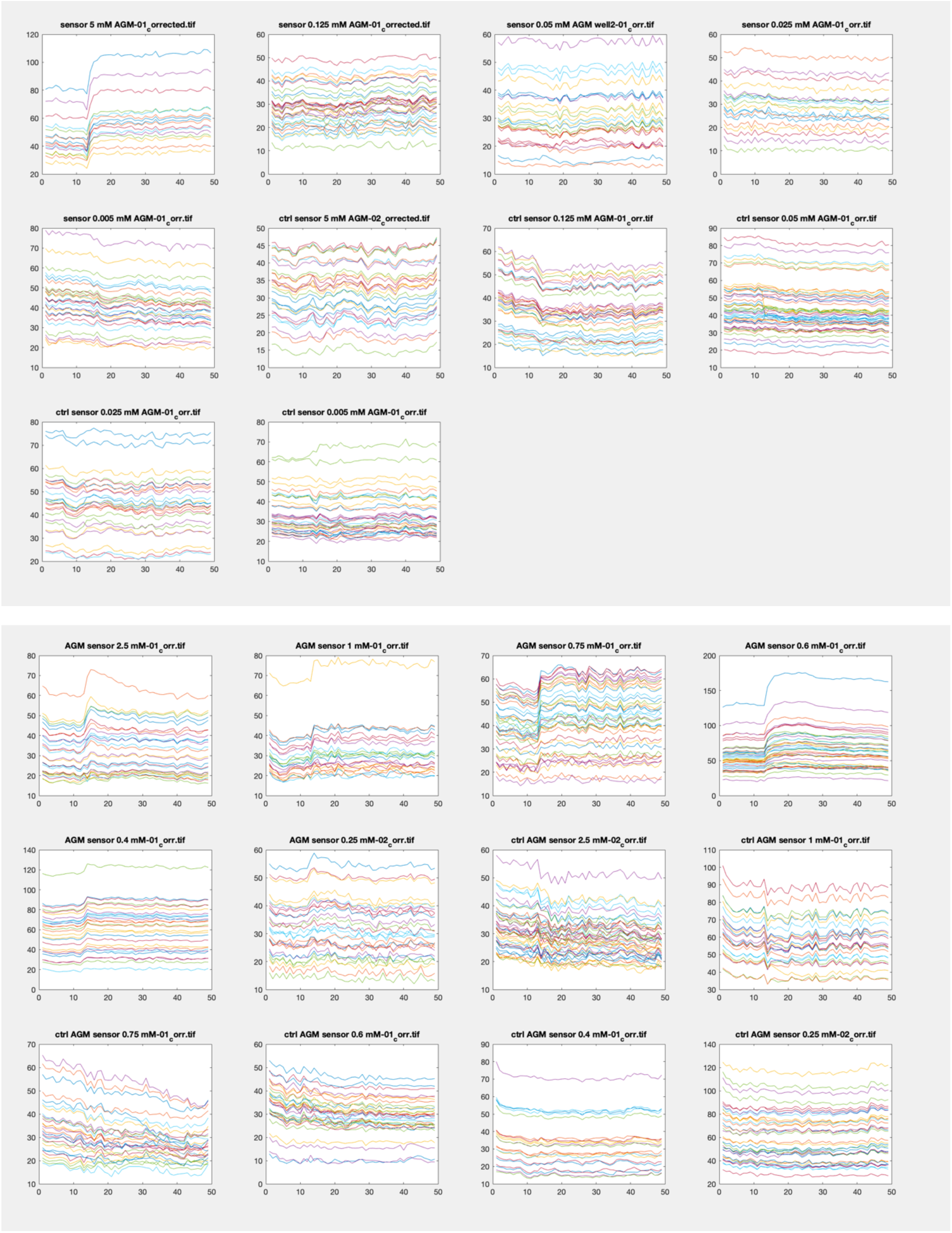
Time traces of individual ROIs. Type of experiment indicated on top. X axis is frame number. Y axis is fluorescence intensity (a.u.).

**Table S1:**
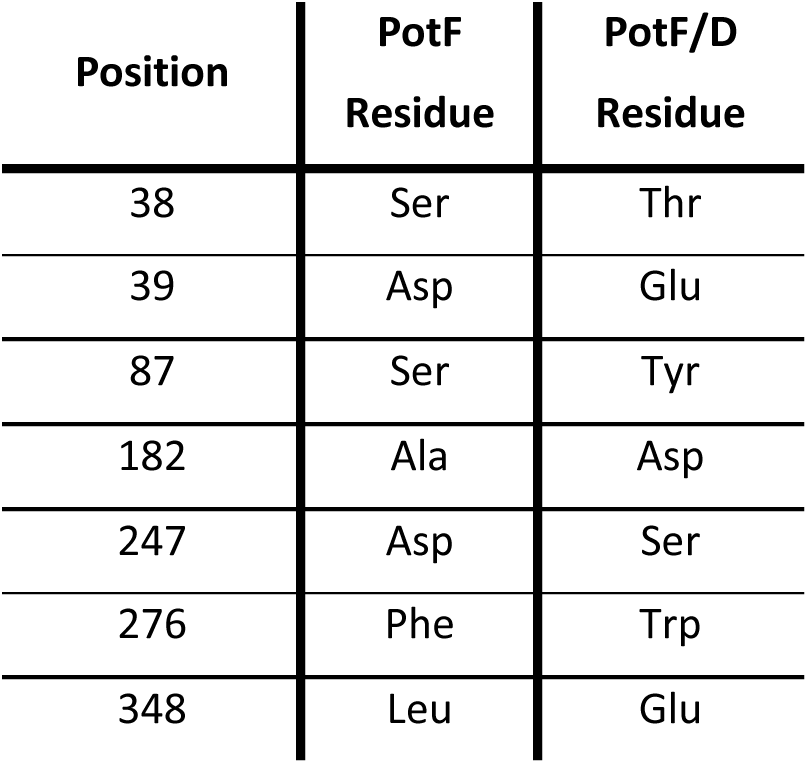
Differences in sequence between PotF and PotF/D.

| **Position** | **PotF**<br>**Residue** | **PotF/D**<br>**Residue** |
| --- | --- | --- |
| 38 | Ser | Thr |
| 39 | Asp | Glu |
| 87 | Ser | Tyr |
| 182 | Ala | Asp |
| 247 | Asp | Ser |
| 276 | Phe | Trp |
| 348 | Leu | Glu |

**Table S2:** ITC data for all measurements. N/D = Not detectable.

| Variant | Ligand | K<sub>D</sub> [μM] | ΔG [kcal/mol] | ΔH [kcal/mol] | -TΔS [kcal/mol] | n |
| --- | --- | --- | --- | --- | --- | --- |
| PotF/D | AGM | 4.0 | -7.2 | -10.3 | 3.1 | 0.90 |
| PotF-S87Y-D247S | AGM | 1.2 | -7.9 | -10.4 | 2.5 | 0.95 |
|  | PUT | N/D | - | - | - | - |
|  | SPD | N/D | - | - | - | - |
|  | CDV | N/D | - | - | - | - |
| PotF-S87Y-A182D | AGM | 0.3 | -8.7 | -5.8 | -2.9 | 1.0 |
|  | PUT | N/D | - | - | - | - |
|  | SPD | N/D | - | - | - | - |
|  | CDV | N/D | - | - | - | - |
| PotF-D247K | AGM | N/D | - | - | - | - |
|  | PUT | N/D | - | - | - | - |
|  | SPD | N/D | - | - | - | - |
|  | CDV | N/D | - | - | - | - |

**Table S3:**
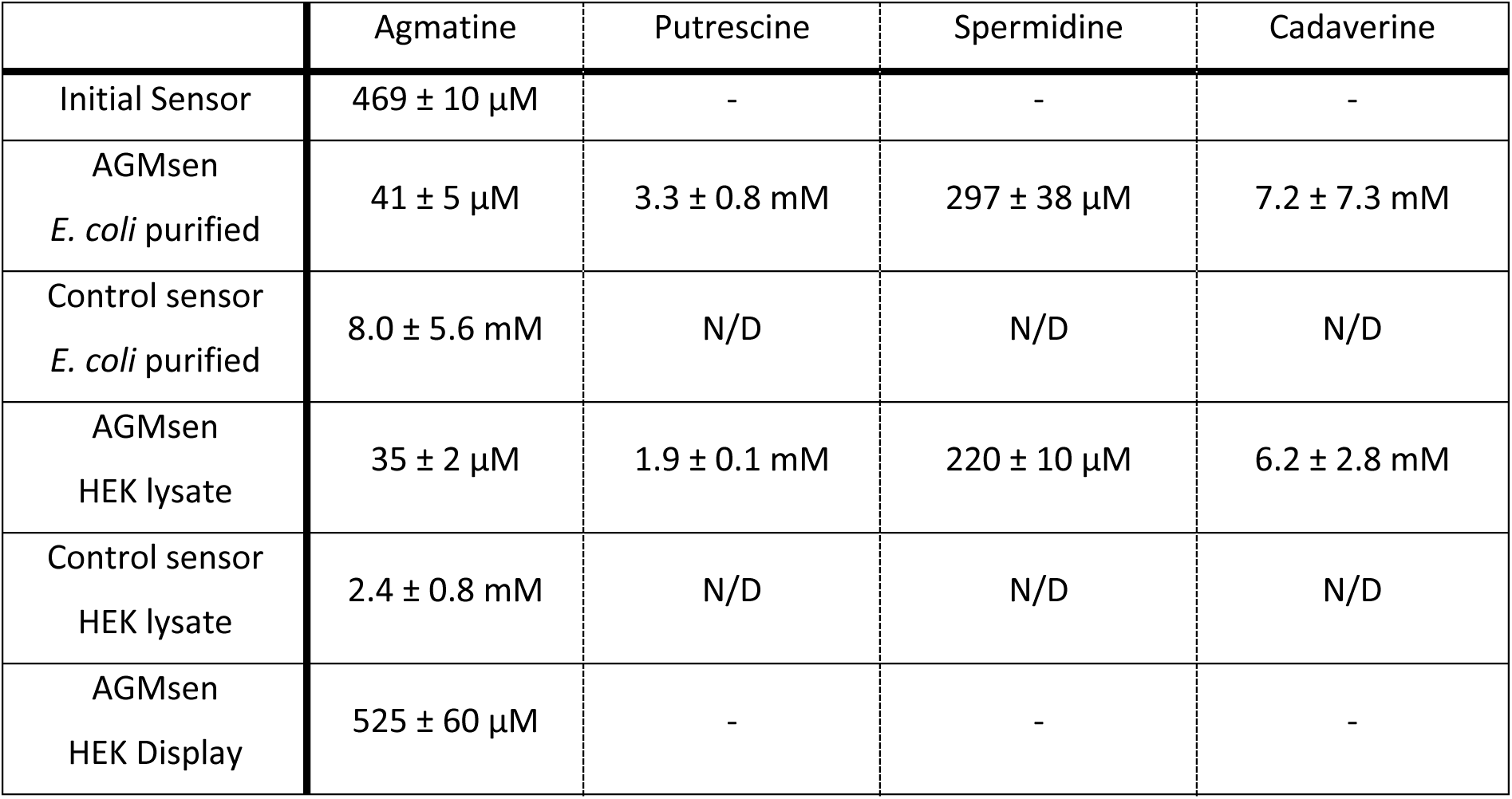
Affinity from dose-response measurements. Data was fitted by Hill-equation using the fit-o-mat. ^30^**. N/D = Not detectable.**

**Table S4:** Crystallographic data and refinement statistics for all solved crystal structures.

|  | PotF-S87Y-A182D | PotF-D247K |  | PotF-S87Y-A182D | PotF-D247K |
| --- | --- | --- | --- | --- | --- |
| PDB ID | 8ASZ | 8AT0 | Refinement |  |  |
| Data collection |  |  | Reflections used in refinement | 86267 (8426) | 54717 (5385) |
| Wavelength [Å] | 0.9184 | | Reflections used for $R_{\text{free}}$ | 2099 (205) | 2098 (206) |
| Resolution range [Å] | 37.33 - 1.28 (1.33 - 1.28) | 45.55 - 2.00 (2.07 - 2.00) | $R_{\text{work}}$ | 0.179 (0.391) | 0.201 (0.306) |
| Space group | P 21 21 21 | P 32 2 1 | $R_{\text{free}}$ | 0.209 (0.380) | 0.243 (0.359) |
| Cell parameter | | | $CC_{\text{work}}$ | 0.970 (0.604) | 0.968 (0.587) |
| a, b, c, [Å] | 37.3 79.1 113.3 | 70.7 70.7 272.6 | $CC_{\text{free}}$ | 0.970 (0.676) | 0.953 (0.460) |
| $\alpha, \beta, \gamma$ [°] | 90 90 90 | 90 90 120 | Number of non-hydrogen atoms | 3305 | 5840 |
| Total reflections | 628916 (62355) | 602533 (61804) | macromolecules | 2851 | 5444 |
| Unique reflections | 86290 (8427) | 54733 (5385) | solvent | 378 | 308 |
| Multiplicity | 7.3 (7.4) | 11.0 (11.5) | Protein residues | 343 | 682 |
| Completeness [%] | 99.0 (98.0) | 99.9 (100.0) | RMS bond lengths [Å] | 0.016 | 0.003 |
| Mean I/sigma [I] | 14.6 (0.6) | 11.2 (0.7) | RMS bond angles [°] | 1.04 | 0.57 |
| Wilson B-factor | 17.9 | 39.34 | Ramachandran favored [%] | 97.1 | 97.4 |
| No. of molecules per a.u. | 1 | 2 | Ramachandran allowed [%] | 2.9 | 2.7 |
| Matthews coefficient |  |  | Ramachandran outliers [%] | 0.0 | 0.0 |
| $R_{\text{merge}}$ | 0.065 (2.904) | 0.181 (2.980) | Rotamer outliers [%] | 1.28 | 0.7 |
| $R_{\text{meas}}$ | 0.070 (3.118) | 0.190 (3.120) | Clashscore | 4.08 | 4.63 |
| $R_{\text{pim}}$ | 0.026 (1.122) | 0.057 (0.918) | Average B-factor | 30.53 | 45.95 |
| $CC_{1/2}$ | 1.000 (0.280) | 0.999 (0.256) | macromolecules | 29.23 | 45.53 |
| $CC^*$ | 1.000 (0.661) | 1.000 (0.639) | solvent | 37.71 | 47.96 |
|  |  |  | Number of TLS groups | 1 | 2 |

**Table S5:**
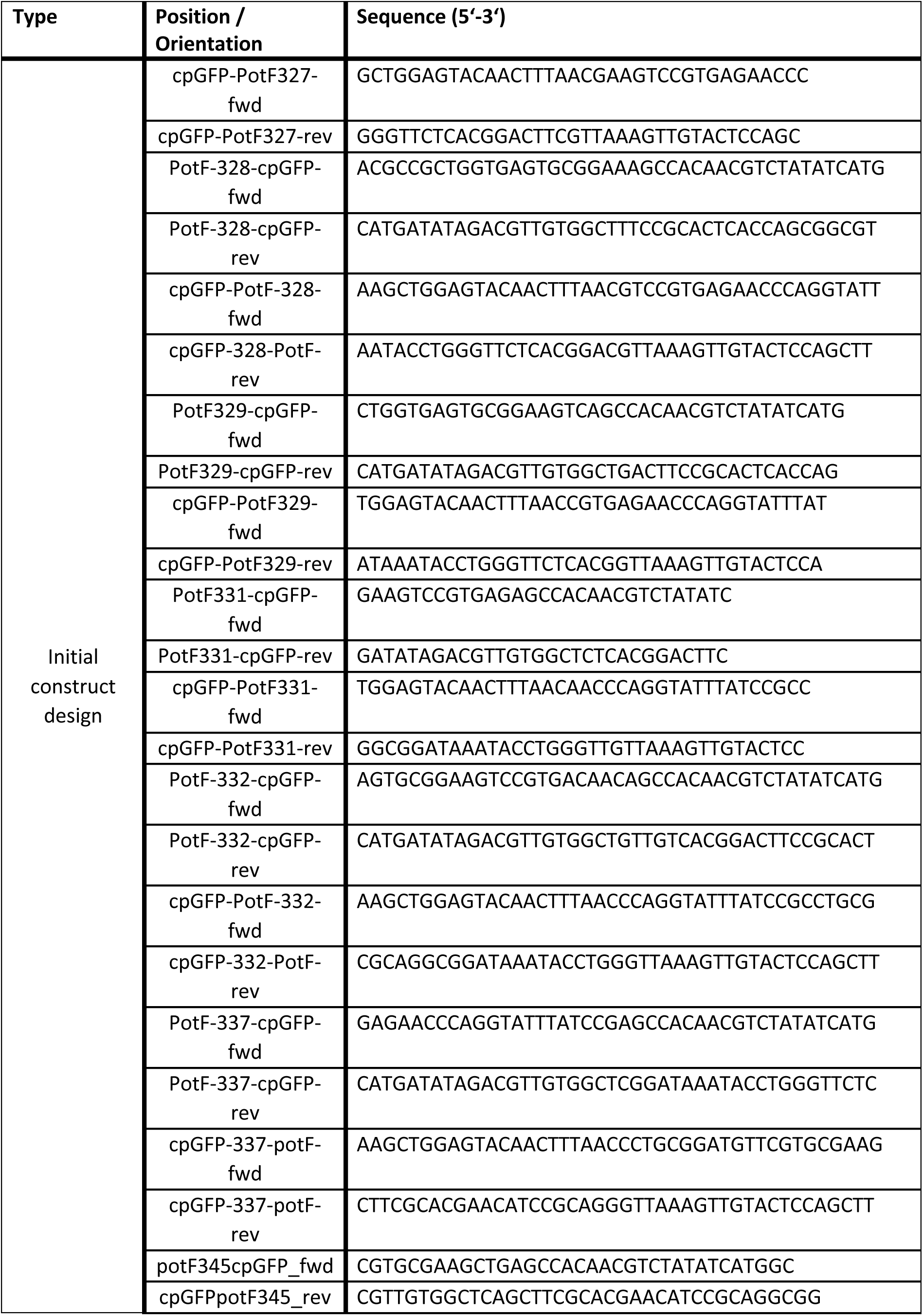
Oligonucleotides used for initial construction of the sensor.

**Table S6:**
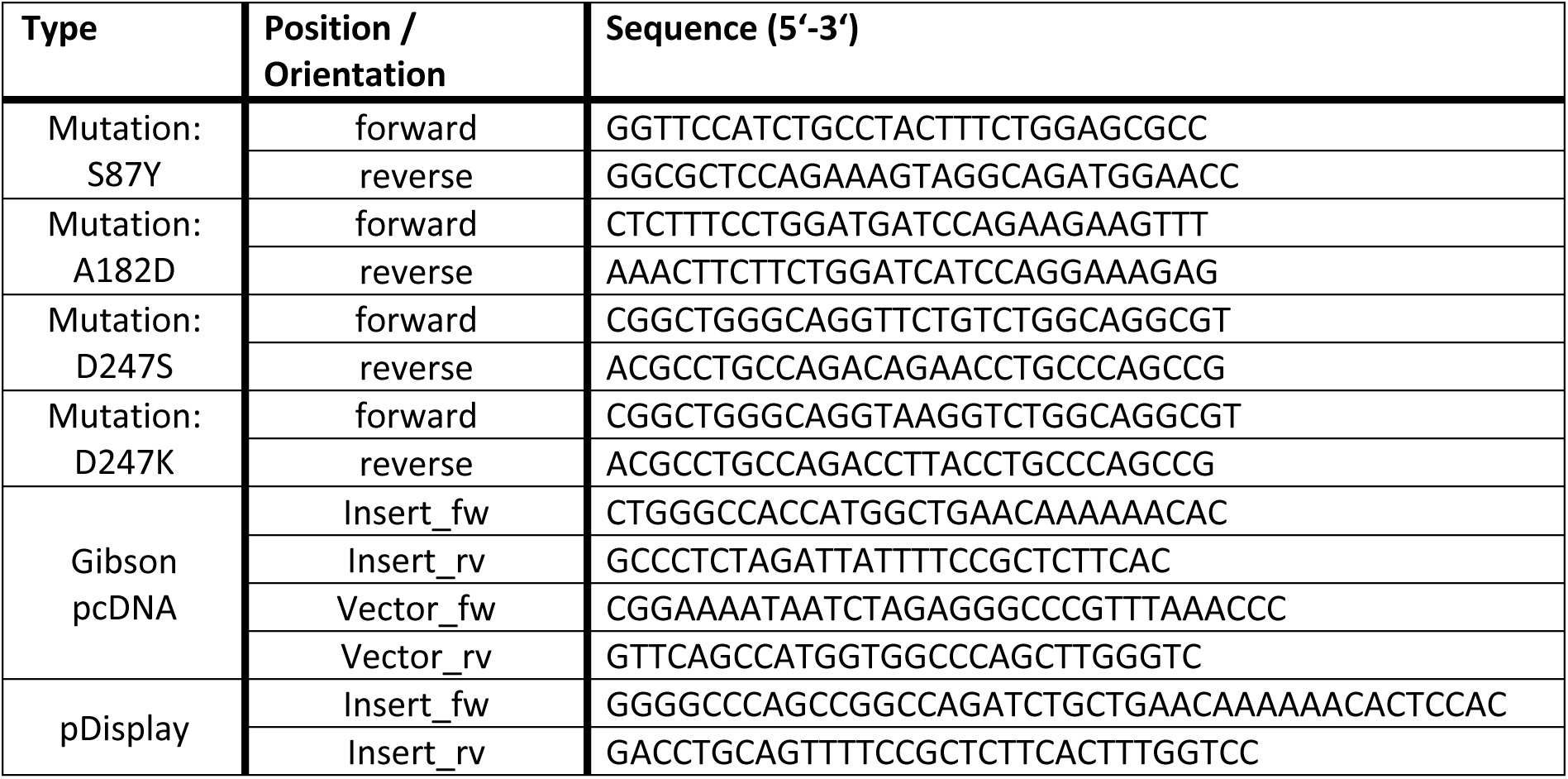
Oligonucleotides used for cloning and the generation of sensor variants via QuickChange.

**Table S7:**
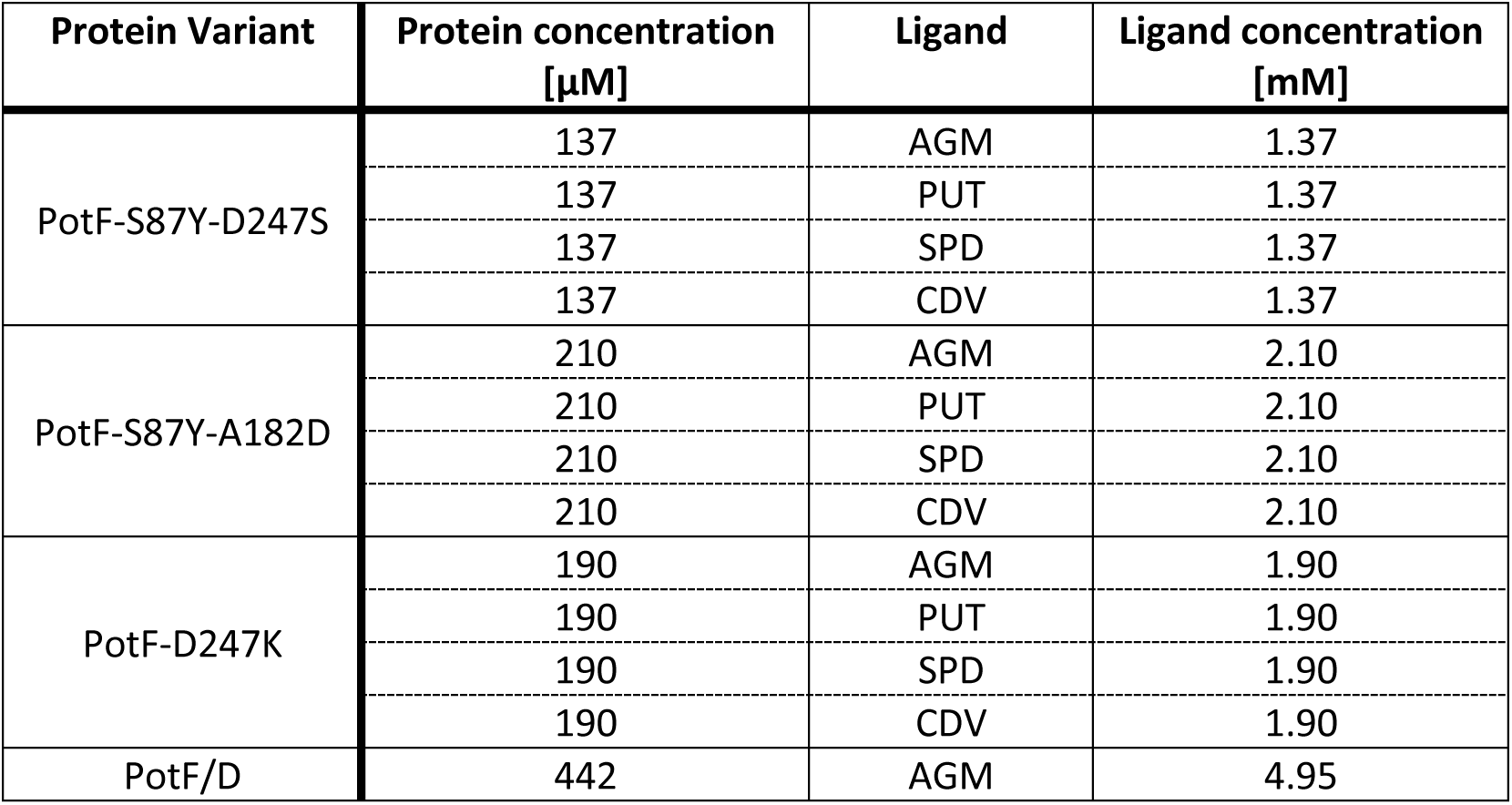
Concentrations of protein and ligand solutions used for ITC measurements.

| Protein Variant | Protein concentration [μM] | Ligand | Ligand concentration [mM] |
| --- | --- | --- | --- |
| PotF-S87Y-D247S | 137 | AGM | 1.37 |
|  | 137 | PUT | 1.37 |
|  | 137 | SPD | 1.37 |
|  | 137 | CDV | 1.37 |
| PotF-S87Y-A182D | 210 | AGM | 2.10 |
|  | 210 | PUT | 2.10 |
|  | 210 | SPD | 2.10 |
|  | 210 | CDV | 2.10 |
| PotF-D247K | 190 | AGM | 1.90 |
|  | 190 | PUT | 1.90 |
|  | 190 | SPD | 1.90 |
|  | 190 | CDV | 1.90 |
| PotF/D | 442 | AGM | 4.95 |

